# Mammals achieve common neural coverage of visual scenes using distinct sampling behaviors

**DOI:** 10.1101/2023.03.20.533210

**Authors:** Jason M. Samonds, Martin Szinte, Carrie Barr, Anna Montagnini, Guillaume S Masson, Nicholas J. Priebe

## Abstract

Most vertebrates use head and eye movements to quickly change gaze orientation and sample different portions of the environment with periods of stable fixation. Visual information must be integrated across several fixations to construct a more complete perspective of the visual environment. In concert with this sampling strategy, neurons adapt to unchanging input to conserve energy and ensure that only novel information from each fixation is processed. We demonstrate how adaptation recovery times and saccade properties interact, and thus shape spatiotemporal tradeoffs observed in the motor and visual systems of different species. These tradeoffs predict that in order to achieve similar visual coverage over time, animals with smaller receptive field sizes require faster saccade rates. Indeed, we find comparable sampling of the visual environment by neuronal populations across mammals when integrating measurements of saccadic behavior with receptive field sizes and V1 neuronal density. We propose that these mammals share a common statistically driven strategy of maintaining coverage of their visual environment over time calibrated to their respective visual system characteristics.

## Introduction

A large body of work clearly shows that we move our eyes to align the central portion of our visual field on goal-relevant objects^1–5^, but even when we are not engaged in a task or even when we are fixating on an object, we incessantly continue to make saccadic eye movements several times a second. Outside of highly controlled experimental setups^4^, the target of many of our gaze changes are not predictable^6^. Similar to humans, many other vertebrates also view the world over a sequence of discrete stable fixations generated from coordinated head and eye movements^7–9^. The distributions of size and frequency of the gaze changes between fixations vary substantially for different mammals, including different non-human primates^10^. Even during passive viewing and irrespective of the presence of a fovea, gaze changes are informative by overcoming inhomogeneity in the sensory representation of different animals and providing receptive fields a unique and updated perspective at each fixation (Fig. 1a, b). The retina in several animals contains a differential density of photoreceptors^11–14^ and occlusion by retinal vasculature^15^. This inhomogeneity persists along visual pathways and is related to cortical magnification factor^16,17^, acuity^18–23^, color sensitivity^24^, and irregular ocular dominance domains^25^. Additional inhomogeneities emerge within cortex, such as the irregular distribution of orientation representation in mice^23^ and disparity selectivity in mice and non-human primates^26–28^. By making gaze changes to cover different regions of the scene, an internal representation may be constructed by integrating novel receptive field information over successive fixations^29–34^.

**Figure 1.**
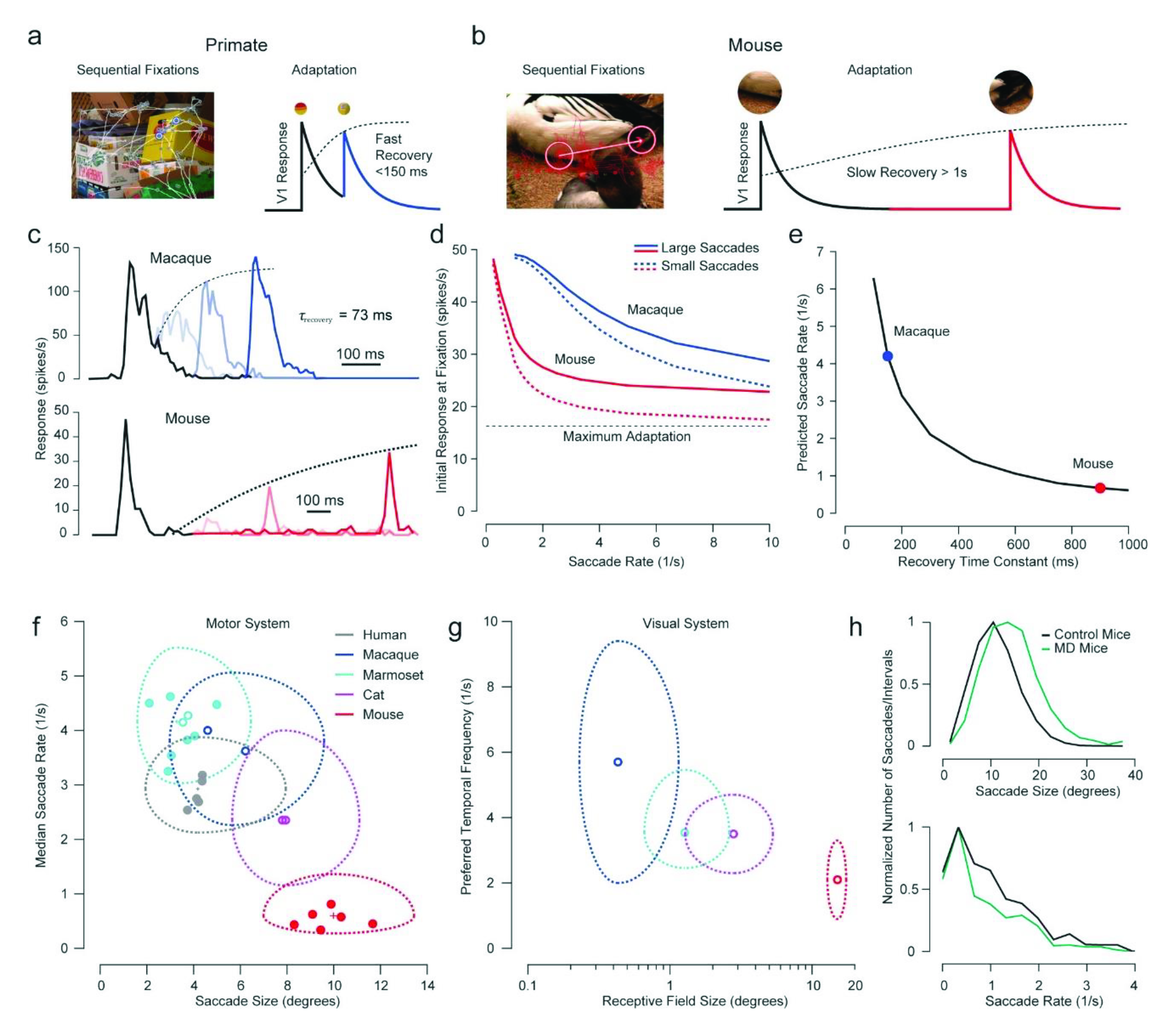
Saccade rates increase with decreasing saccade size matching faster recovery from adaptation for smaller receptive fields. **a**, **b**, Sequential fixations change receptive field inputs and differences in adaptation recovery time constants allow primates (**a**) to make more frequent saccades compared to mice (**b**). **c**, Top row: example macaque V1 responses to two repeated presentations of the same stimulus^38^. Black represents neural activity to the first presentation and different shades of blue depict the neural activity after the second presentation in progressively more delayed conditions. Bottom row: example mouse V1 responses to a similar experiment^41^. See model examples in Supplemental Fig. 1. **d**, Magnitude of initial model responses for different saccade rates for macaques and mice. For large saccades, the stimulus changed randomly after a saccade, while for small saccades, the stimulus stayed the same after a saccade. **e**, Saccade rates for a range of recovery time constants that maintain 75% of initial model responses. **f**, Saccade rate versus size distributions for five species. Each solid point represents the medians of a single subject. Open circles of marmosets^43^, macaques^43^, and cats^100,101^ represent previously published data from other laboratories (see Methods). The large dashed outline represents the central half of the distribution (between the 25th and 75th percentiles) and the small cross in the center represents the standard error of the median for the distribution of all subjects. Due to the large number of samples, the standard error bars are smaller than even the size of the data points. See full distributions in Supplemental Fig. 2. **g**, Preferred temporal frequency versus receptive field size distributions from previously published data (see Methods)^18–21,49–59^. The open circle represents the median and the large dashed outline represents the central half of the distribution (between the 25th and 75th percentiles). **h**, Saccade size increases^10^ and saccade rate decreases in mice with monocular deprivation during the critical period (N = 4 mice: n = 1448 saccades) compared to control mice (N = 3 mice: n = 1586 saccades).

Our hypothesis is that neural adaptation and gaze changes work together to provide novel receptive field inputs and conserve energy. Neurons quickly adapt their responsiveness to unchanging visual input^35^, which should lead to the conservation of metabolic energy by reducing overall neural activity^36,37^. Eye and head movements, which change where a receptive field samples, could be continuous and uniform across the visual scene, but that would be inefficient and prevent sufficient processing that requires a stable view of images^9^. To conserve energy and allow the nervous system to process visual inputs, gaze changes are discrete and quick only moving on average the minimum distance necessary to provide novel inputs to the receptive fields given natural scene statistics^10^. This argument successfully accounts for the differences in the distribution of saccade amplitudes across mammals^10^, but it is not clear if it also explains differences in saccade rates. Since sampling the visual environment depends on both space and time, we expanded our conceptual model to include adaptation dynamics, examined the differences in visual response properties and spatiotemporal saccade statistics across different mammals in new and previously published data, and measured visual coverage across time in these animals.

## Results

The adaptation properties of cortical neurons are remarkably distinct across mammals. For both mice and macaques, the responses to brief presentations (<0.5 s) within visual receptive fields decay within a couple hundred milliseconds, but macaque neurons recover their responsiveness very quickly (within a few hundred milliseconds, Fig. 1a, c, blue)^35,38,39^ while mouse neurons recover much slower (over several seconds, Fig. 1b, c, red)^40,41^. Recovery time constants have not been explicitly measured in cat neurons, but the available data suggests that they recover with a time constant somewhere in between these two values^42^. Recovery to adaptation for these animals in these studies were measured for repeated presentations of the same stimuli while the animal was fixating. Saccadic eye movements that decorrelate input across fixations may decrease the effects of neural adaptation^10^, but the adaptation that persists nonetheless requires time for recovery^39^. Macaque neurons will still recover much faster than mouse neurons even with large enough saccades.

To quantify how different saccade rates influence adaptation, we constructed a model in which the dynamics of adaptation could be incorporated into the theoretical framework we previously used to explain the relationship between neural adaptation and saccade size^10^. Responses (*r_m_*) of a population of *M* = 500 neurons to *N* = 500 stimuli (*s_n_*) following sequential saccades were multiplied by a divisive change in gain dependent on the response to the previous stimulus (*s_n_*_-1_):

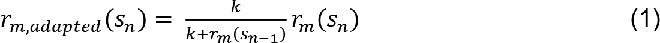

Parameters of this function (*k* and average responses) were chosen to approximately match observed adaptive changes in gain in macaques, mice, and cats, where the average gain never reaches zero for even the fastest repeat presentations (see Supplemental Fig. 1 and Methods for details)^38–42^. We varied the time between stimuli to represent different saccade rates and model responses decayed over time (*t*) as an exponential decay function with a fixed time constant *τ*_*decay*_. The divisive gain based on the initial response after each saccade recovered over time based on a second exponential decay function:

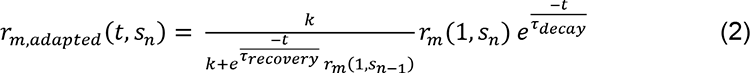

We varied the time constant (*τ_recovery_*) of this function to cover a range that includes experimentally observed adaptation recovery time constants to brief presentations of stimuli in macaques and mice^38–41^. To mimic the effects of saccades, we repeated this analysis but changed the stimulus. For some conditions, the stimulus changed randomly, thus providing a decorrelated input mimicking large saccades. For other conditions, the stimulus did not change, mimicking small saccades relative to receptive field sizes. As saccade rate increases, neurons have less time to recover from adaptation and the overall responses decline (Fig. 1d). For both animals, large saccades reduced adaptation compared to small saccades yielding comparatively higher responses for faster saccade rates, reflecting a linkage between saccade size and rate (Fig. 1d, solid vs. dashed lines). To illustrate how the recovery time would relate to saccade rate, we plotted the saccade rate that would still maintain 75% of responsiveness against the recovery time constant (Fig. 1e). This simple model predicts a dramatic difference in saccade rate that depends on the adaptive dynamics in visual neurons, and captures the large differences in saccade rates observed between macaques (Fig. 1e, blue point; 3.8 saccades/s)^43^ and mice (Fig. 1e, red point; 0.6 saccades/s)^10^.

Our model suggests that there should be a functional link between the spatiotemporal dynamics of the visual and motor systems. To characterize the spatiotemporal tradeoffs made by these two systems, we first recorded and compared the saccade rates and sizes when presenting natural scenes to three different species known to have different visual system properties: humans, marmosets, and mice (Fig. 1f). As demonstrated previously^10^, there are clear differences in saccade sizes between these species, with all animals exhibiting a skewed distribution in saccade amplitude (Supplemental Fig. 2b). Marmosets have the smallest saccades and mice have the largest ones. For all animals, saccade sizes were slightly larger for larger images (see also ref. 44) (Supplemental Fig. 2b; bootstrapped, p < 0.001 for all comparisons). There are also clear differences in saccade rates between species (Fig. 1f). Marmosets have the highest saccade rates and mice have the lowest saccade rates. For all animals, saccade rates were slightly higher for larger images (see also ref. 44) (Supplemental Fig. 2c; bootstrapped, p < 0.001 for all comparisons). Saccade behavior is similar between animals with only a shift in rates and sizes. Including previously reported data from other animals (open circles for marmosets, macaques, and cats) reveals an inverse relationship between saccade rate and saccade size across distinct mammalian species (Fig. 1f). The saccade size and rate statistics reported in several publications are remarkably similar to our data when the authors employed similar “free viewing” approaches and image size and duration are considered^44–48^. Humans and all animals were passively viewing natural images of the same or similar size as humans and were prevented from making head movements. One distinction was that mice were allowed to freely run, but their data were similar whether they were running or stationary (except a lower rate when stationary; see ref. 10).

The decorrelation hypothesis predicts that saccade size differences between species are attributed to the properties of their respective visual systems^10^. Human and macaque receptive fields are small relative to mouse receptive fields, whereas marmoset and cat receptive fields lie between these sizes. Consequently, to achieve decorrelation and sample new information, mice need much larger saccades in order to move their larger receptive fields as compared to humans and the other species. Our model also suggests that saccade rates will be matched to differences in adaptation recovery time constants (Fig 1e). Since adaptation time constants in individual animals can vary depending on experimental conditions and data based on similar conditions is fairly limited across species, we rather considered preferred temporal frequency distributions across several species to provide a more comprehensive view of the relationship between spatial and temporal properties of their visual system neurons. Indeed, temporal frequency tuning has been measured in several animals under very similar experimental conditions (e.g., spike rate responses to sinusoidal luminance gratings; see Methods) in different laboratories^49–59^. Similar to the saccade statistics, there is an inverse relationship between preferred temporal frequency and receptive field size across these animals (Fig. 1g). Both the oculomotor and visual systems share spatiotemporal tradeoffs across species where those with smaller receptive fields can make smaller saccades, but require more frequent sampling and faster visual processing. Indeed, we find that low luminance contrast of natural images, which it is known to shift temporal frequency tuning lower^54,60,61^, is associated with a corresponding reduction in saccade rate (Supplemental Fig. 3).

The relationship we uncover between saccade size and rate may result from differences between species other than receptive field sizes. To test whether changes in receptive field size are linked to saccade rate, we reduced the spatial acuity of mice using monocular deprivation during the critical period (age 24-32 days). This manipulation shifted their saccades to larger sizes (Fig 1h, top; see ref. 10) and significantly reduced their saccade rates compared to control mice (Fig 1h, bottom; bootstrapped, P < 0.001). This causal manipulation demonstrates that the inverse relationship found between saccade size and rate is related to visual functional properties such as acuity, and presumably receptive field sizes. A similar change in saccade size and rate is observed in humans on a coarse scale with substantial reduced acuity due to macular degeneration or foveal occlusion^62,63^ and on a finer scale in subjects with myopia and amblyopia^64,65^.

Our modeling and empirical data in different species show that visual and motor systems share similar spatiotemporal tradeoffs to sample visual information. We reasoned that such a tradeoff is constrained by a common computational goal: achieving equivalent novel (decorrelated) spatiotemporal coverage of their visual environment. The mouse with both large V1 receptive fields and eye movements would need to make eye movements less frequently compared to the human with both small receptive fields and eye movements in order to achieve similar updated coverage over time.

To assess coverage over time quantitatively, we applied a circular mask equal to the median V1 receptive field size for each animal to every fixation point (Fig. 2a, green circles). We then added up all of those masks (each equal to one for that area) across space and time for all images to generate cumulative spatial maps. Lastly, we divided those maps by the total of the intersaccadic interval times to get a percent coverage per second (Fig. 2a, right). Because coverage is based on area, the mouse achieves much more coverage over time compared to marmosets and humans even with their slower saccade rates (Fig. 2b).

**Figure 2.**
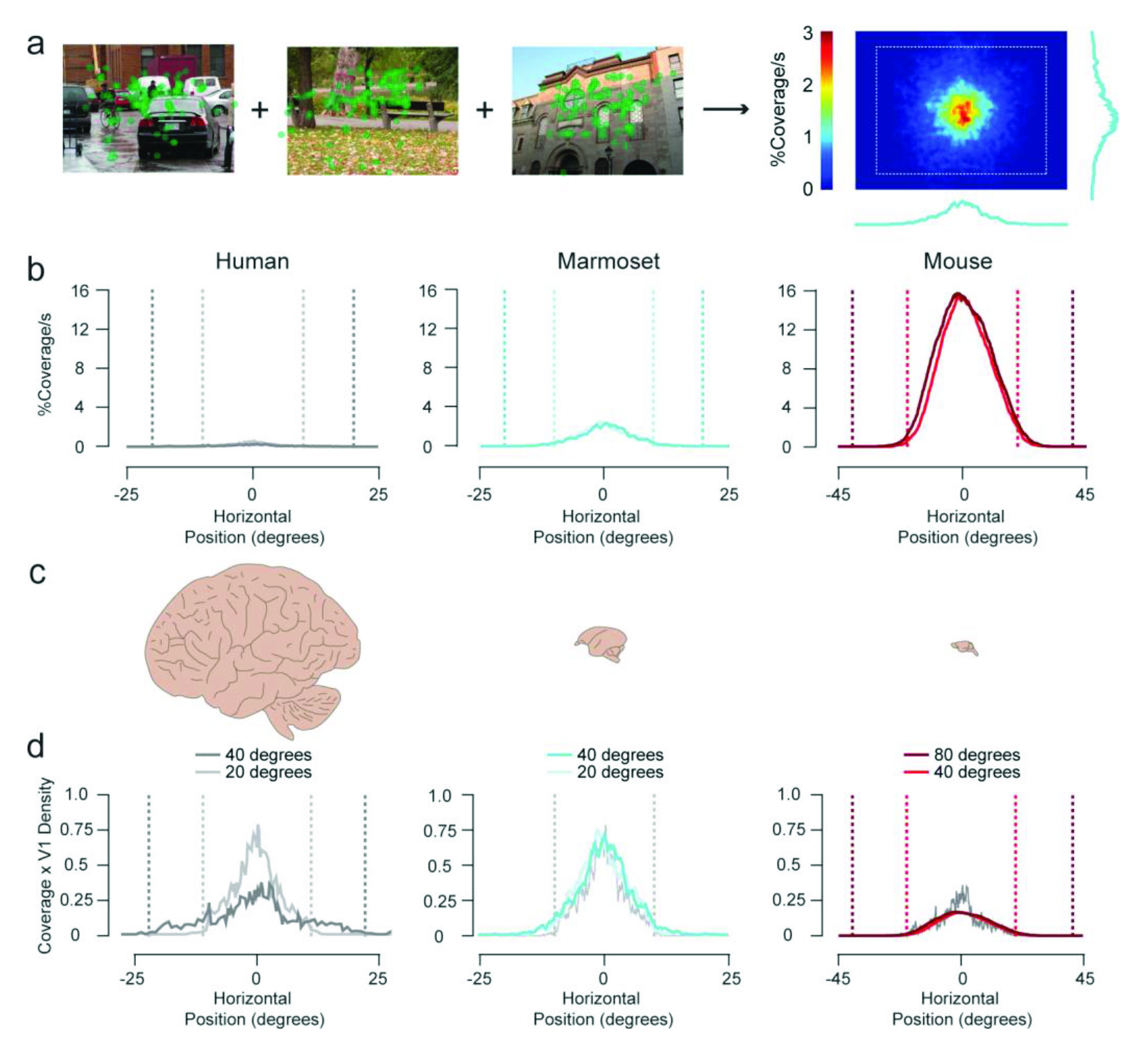
Fixations over time provide similar visual processing coverage across different mammals. **a**, Computing average receptive field coverage over time from saccades. **b**, Horizontal cross sections of receptive field coverage over time for each mammal. Lighter colors are data obtained when subjects were viewing small images. Vertical dashed lines represent image boundaries. **c**, Relative brain sizes between each mammal. **d**, Horizontal coverage data from **b** multiplied by the average two-dimensional V1 cell density for each mammal. Human data are replotted with marmoset and mouse data to facilitate comparison.

One notable exception to the general inverse relationship between saccade rate and size across species is observed for humans, who appear to have lower saccade rates than macaques although they have slightly smaller saccade sizes (Fig. 1f). In addition, if we assume that marmosets make larger saccades with their heads^43^ with no decrease in saccade rate (see Supplemental videos 1-2 and Supplemental Figs 4-5), they would have higher saccade rates than macaques with larger saccade sizes. This suggests that saccade rates might also depend on species-specific factors other than saccade sizes and receptive field sizes.

We postulated that the main bottleneck constraining coverage over time is the receptive field size and density of the first cortical stage, the primary visual cortex. The animals we are examining have very different densities of V1 cortical neurons per degree of visual angle (dva), and therefore differences in densities of feature preferences^66^ (Fig. 2c). Humans have many more neurons for each receptive field location representing many more different visual features than mice and mathematically, larger networks can provide greater decorrelation^67^. We can take our original coverage maps over time (Fig. 2b) and multiply by the two-dimensional (2D) V1 neuronal density for each animal (see Methods for details) to compare receptive field cortical coverage over time between species (Fig. 2d). Because marmosets often use head movements for gaze changes and their eye positions, unlike humans, appear to be restricted to mostly ±10 dva (Fig. 2b, center; see also Supplemental videos 1-2 and Supplemental Figs 4-5 and ref 43), we observe comparable coverage for smaller 20 dva images between humans and marmosets (Fig. 2d, center, light gray versus light cyan). Mice also use head movements for changes in gaze^68–70^, but their eye positions extend out to at least cover ±20 dva (Fig. 2b, right) so we observe comparable coverage for 40-dva images between humans and mice (Fig. 2d, right, dark gray versus red). These comparisons suggest that saccades for these mammals are decorrelating similar numbers of neurons over time despite having very different saccade rates, saccade sizes, and receptive field sizes. With this sampling strategy, differences in saccade rates and sizes between animals are explained by corresponding differences in receptive field sizes and cortical densities with respect to visual space. If saccade rates and sizes are calibrated to achieve a certain level of visual coverage, this would also explain why saccade rates and sizes change with image size and decrease over viewing time (Supplemental Figs. 2 and 6; see also refs 10,44). As our model predicts, saccade rates are always higher in the condition where it is easier to achieve decorrelation. The oculomotor system indeed seems to favor larger versus smaller saccades, since it is easier and faster to make larger saccades^71–73^.

## Discussion

We propose that gaze changes efficiently sample visual scenes based on natural statistics, neural receptive fields, and adaptation, but this hypothesis makes some important assumptions. First, for the purposes of predicting saccade sizes, we assumed that the receptive field is static and only examined image differences across sequential fixations^10^. In fact, saccades have complex dynamics over 10’s of milliseconds and the eyes drift and oscillate continuously over 100’s of milliseconds or seconds of fixation. These fast and slow oculomotor dynamics effectively whiten the visual inputs^74–76^. This whitening is typically outside of the frequency spectrum of the V1 response preferences that we used to predict saccade sizes (Supplemental Fig. 7). Indeed, V1 responses following a saccade are very similar to those evoked by sequentially flashed stimuli^77^. Nonetheless, saccades can produce an intra-saccadic flow of information that is sensed^78,79^, can boost post-saccadic visual motion processing^80^, and have functional consequences for spatial vision^81,82^. Second, we assumed that the primary visual cortex was the main processing bottleneck that constrains saccade sizes and rates. This choice was partly made because of the large number of comparable data available across multiple species. This assumption does not rule out that differences in receptive fields found earlier, later, or subcortically in the visual system between species contribute to their oculomotor differences. Future work shall include additional processing that occurs during both saccades and fixation and how they influence both cortical and subcortical responses. Another possibility that we did not address in this work is that the oculomotor spatial and temporal species-specific properties have actually contributed to shape the properties of the visual system rather than the oculomotor behavior being the consequence of the visual system characteristics^83^.

Yarbus illustrated that humans move their eyes deliberately towards objects and features of interest and the overall pattern of those movements can heavily depend on the task^1^. Decades of research support that the primary purpose of saccadic eye movements is to move an area of interest into the high-spatial acuity portion of the visual field^1–5^. Even animals with no fovea, such as mice, use gaze changes to move objects of interest into the central portion of their upper visual field^69,84^ where binocularity and spatial features are represented better^22,23,27^. Our hypothesis is complementary to foveation and does not attempt to predict saccade targets instantaneously. We use decorrelation as a measurement to quantify a statistical sampling strategy for natural scenes over a long time scale of several fixations to update an internal visual representation. It is important to point out that this sampling strategy can depend on and adapt to environmental and task demands as well. We see this when varying image contrast and size (Supplemental Figs. 3 and 6; see also ref. 44), but previous studies have also shown predictable changes in coverage (size/rate) statistics using images with biased spatial frequency content^85^, tasks with high acuity demands^86^, or tasks with high cognitive demands^45,48^. Previous work examining transsaccadic integration also shows that information from past fixations influence perception during the current fixation. If that past information about a particular visual feature was more reliable than the current information, it has a stronger influence on perception^32,33,87,88^. Finally, a higher rate of saccades improves performance in detecting changes within a scene^89^ highlighting the importance of maintaining sufficient coverage. Overall, our analyses illustrate that the oculomotor and visual systems of multiple mammals coordinate to sample the environment efficiently under a diverse range of processing constraints.

## Materials and Methods

### Adaptation model

For the model, we chose parameters for equation 2 that best matched the average response dynamics (decaying to 30% of the initial response) to brief sequential presentations reported for adaptation experiments in macaques and mice^38–42^. The decay time constant *τ*_*decay*_ for all simulations was 60 ms and the recovery time constant *τ*_*recovery*_ varied between 100 and 900 ms. The initial non-adapted firing rate was 50 spikes/s and the constant *k* for the divisive gain function was 30.

### Human participants

Five participants of the Institute des Neurosciences de la Timone (22-29 years old, 2 females) took part in the free viewing tasks. Experiments were approved by the Ethics Committee of Aix-Marseille University and conducted in accordance with the Declaration of Helsinki. All participants gave written informed consent.

### Mice

Experiments and procedures were performed on eight C57bl6 adult female and male mice (RRID: JAX: 00664), one female and two male PV-Cre mice^90^, two female PV-Cre;Ai14 mice^90^, and two female and four male PV-Cre;ChR2 mice^91^ (all from the C57bl6 line of mice). Mice were group housed and maintained on a 12-hour light/dark cycle under standard housing conditions. All procedures and care were performed in accordance with the guidelines of the Institutional Animal Care and Use Committees at the University of Texas at Austin.

### Marmosets

Seven marmosets (1.5-4 years, 2 females) were obtained from TxBiomed and bred in-house and were group-housed. Food and water were provided ad libitum. All procedures and care were performed in accordance with the guidelines of the Institutional Animal Care and Use Committees at the University of Texas at Austin.

### Human saccades

For free viewing experiments, participants sat in a quiet and dimly illuminated room, with their head positioned on a chin and forehead rest. The experiments were controlled by a HP Z420 (Hewlett-Packard, Palo Alto, CA, USA) computer equipped with an Nvidia Quadro 600 video card (Nvidia, Santa Clara, CA, USA). Binocular eye position was recorded using an Eyelink 1000 tower mount (SR Research, Osgoode, ON, Canada) at a sampling rate of 2 kHz (1 kHz per eye). The experimental software controlling the display as well as the eye tracking was implemented in Matlab (The MathWorks, Natick, MA, USA), using the Psychophysics^92,93^ and Eyelink toolboxes^94^. Stimuli were presented at a viewing distance of 60 cm on a 32-inch Display++ LCD monitor (Cambridge Research System, Rochester, UK) with a spatial resolution of 1920 by 1080 pixels and a vertical refresh rate of 120 Hz. Participants’ gaze position was calibrated with a 13-point custom calibration sequence with sequences validated at the beginning of an experimental session as well as when necessary.

Participants took part in free viewing tasks composed each of 40 runs of eight trials each. These runs were completed in four experimental sessions (on different days) of about 60 min each (including breaks). Participants were instructed to freely explore a set of 80 images displayed at the center of a gray background screen. These pictures were presented on each trial for a duration of 30 seconds and separated each by two seconds of blank gray screen. Pictures were selected from the natural images from the McGill Calibrated Color Image Database^95^. For the first 20 runs, participants randomly explored the screen with either the 80 full-sized selected images covering 40 by 32 dva of the visual field or the same 80 images center-cropped covering 20 by 16 dva of the visual field. For the next 20 runs, each participant explored 40 images selected randomly from the same set of 80 images, but with their luminance contrast scaled down uniformly at either 5 or 10% of the original contrast. These images covered 40 by 32 dva of the visual field during the first 10 runs and 20 by 16 dva of the visual field in the last 10 runs. Data from images of both sizes were combined for contrast-related results.

For all experiments, saccades were detected as previously described for marmosets with a velocity threshold^10^.

### Marmoset saccades

Marmoset eye position was tracked as previously described using an EyeLink 1000 (SR Research, Osgoode, ON, Canada) camera to capture binocular eye position at sampling rate of 1 kHz (500 Hz for each eye)^10,27^ and saccades were detected with a velocity threshold^10^. Twenty-four small (20 by 16 dva) and 24 large (40 by 32 dva) images from the McGill Calibrated Color Image Database^95^ were presented per session on a CRT monitor (85 Hz refresh rate) 50 cm away randomly interleaved them with unrelated 1-s fixation trials using Maestro software suite (https://sites.google.com/a/srscicomp.com/maestro/). In separate sessions, the images were presented with their luminance contrast scaled down uniformly at either 5 or 10% of the original contrast. Data from images of both sizes were combined for contrast-related results.

Marmosets had higher saccade rates than expected with respect to their temporal frequency relationship to the other animals (Fig. 2f-g). Previously, we found that marmosets have smaller saccade sizes than we predicted based on their receptive field sizes^10^. We attributed this mismatch to marmosets normally using saccadic head movements to make rapid changes in gaze^43^. To probe this idea further, head and eye position were measured in marmosets that were not head restrained using DeepLabCut^96^ to track pupils and the center of the upper forehead in 400 by 400 pixel images collected at 60 frames per second (see Supplemental video 1).

Calibration for these results was done on head restrained marmosets using the fixation methods described previously^43,97^ (see Supplemental video 2). Saccade statistics based on DeepLabCut of posted marmosets were not distinguishable from those measured with the EyeLink system (Supplemental Fig 2). For analysis of unposted marmosets, we only looked at continuous segments when the gaze was towards the monitor (within ±50 dva from the monitor center).

Marmosets clearly used combined head and eye movements for nearly all changes in gaze, which always produced larger gaze changes than with the eyes alone (Supplemental Fig 3a-c). We did not observe obvious differences in the gaze change rates (Supplemental Fig 3d).

Interestingly, marmosets also cock their heads frequently and rapidly, especially when presented with novel objects^98^. This might increase their ability to decorrelate orientation tuned receptive field responses and allow them to sample visual information at a faster rate than with saccades alone (shift the curve in Fig. 1d to the right).

For comparison to our data, the two open circles (median size and rate for each subject) in Fig. 2f were reformatted from figures from a previous publication and the two marmosets were viewing 44 by 34 dva images displayed for 20 s of natural images including humans, macaques, and marmosets^43^.

### Mouse saccades

For image size experiments, saccade statistics are from four C57bl6 and two PV-Cre;Ai14 female mice from a previously reported study^10^. These mice were trained for a binocular discrimination task^27^ and 8-12 small (40 by 32 dva) and 8-12 large (80 by 64 dva) images/per session were presented before or after 2-3 training sessions. Images from the McGill Calibrated Color Image Database^95^ were presented onto a screen 22 cm away with a DLP LED projector. Binocular eye movements were tracked using custom software to analyze 250 by 250-pixel images collected at 20-30 frames per second with cameras aligned with the orbital axis of each eye. Since mice have a much higher contrast threshold and saturation point than primates, for image contrast experiments, we had them view natural images scaled down to 25 and 50% of the original contrast. This increase for mice compared to primates roughly corresponds to the ratio of increase in contrast thresholds for mice compared to humans^99^. For three male and one female C57bl6 mice, images were scaled to modulate at a ratio of 0.25 or 0.5 of their original contrast modulation. Experiments were run in the same manner as the previous study except four images each of 25, 50, and 100% contrast were randomly interleaved for two sessions. Mice were first trained to discriminate black from white or crossed versus uncrossed disparities from dynamic random dot stimuli by licking left or right. Natural images were shown before or after a training session when mice reached consistent performance above 75% correct for any task. Eyes were tracked in the same manner as previously described except that DeepLabCut^96^ was used to track pupil position. Saccades were detected as previously described with a velocity threshold and comparing detection for both eyes^10^.

For monocular deprivation (MD) experiments^100^, one eye in each mouse (two female and two male PV-Cre;ChR2) was sutured closed at P24 under anesthesia and sutures were removed at P33. Three age-matched PV-Cre mice (one female and two males) were used as controls. Acuity was assessed and saccades for all mice viewing natural images were measured at P40^10^.

### Macaque saccades

Macaque data in Fig. 3A were reformatted from figures from a previous publication (median size and rate for each subject) and the two macaques were viewing 44 by 34 dva images displayed for 20 s of natural images including humans, macaques, and marmosets^43^. The median and 25th and 75th percentile was also extracted from the aggregate distribution of both subjects.

### Cat saccades

Cat saccade size statistics (median, 25th percentile quartile, and 75th percentile quartile of reported sizes) were reformatted from figures from previous publications where the cats were either free viewing in the dark^101^ or viewing a 21-min natural video of unknown size^102^. Cat saccade rate statistics (median, 25th percentile quartile, and 75th percentile quartile of intersaccadic intervals) were extracted from horizontal and vertical traces of unpublished data of a head-fixed cat watching a 30-s video of animals at the zoo (unknown size, but smaller than the 60 by 40 dva screen) provided by Theodore Weyand at Louisana State University School of Medicine. Saccades were detected as previously described for mice and marmosets with a velocity threshold. Dr. Weyand observed that cats made saccades more often for the movie compared to dots or a blank screen (unpublished observations).

### Receptive field size and temporal frequency data

All receptive field size distributions were extrapolated from data from previously published studies that measured the areas or widths of the minimum response field from electrophysiological recordings for cats, marmosets, and macaques, and GCaMP fluorescence thresholded at half the maximum response for mice. Receptive field (RF) size distributions for cats^18–20^, marmosets^17^, and macaques^17,21^ were generated by combining receptive field size versus eccentricity data fits with cortical magnification data fits, and for mice, we used the reported distribution^103^ (see reference 10 for more details).

All preferred temporal frequency distributions were aggregated from previously published electrophysiology studies that were based on peak responses in tuning curves generated from average spike rates measured from single neurons in anesthetized animals. For all studies, responses for each temporal frequency were measured using drifting sinusoid gratings centered on the receptive field of single neurons using gratings with the preferred orientation and spatial frequency of the neuron. All spikes were measured extracellularly for single units. All studies used single insulated tungsten electrodes, except a silicon microprobe with 16 recording sites was used to record spikes in mice for one study^55^. Only summary statistics from the microprobe study were used to validate our aggregate statistics from other studies. For macaques, peak temporal frequencies for 130 single neurons from ref. 50 were combined with fitted peak temporal frequencies for 75 single neurons from ref. 51 to generate an aggregate distribution of peak temporal frequencies. This distribution was consistent with summary statistics reported in additional macaque studies using similar methods^53,54,57^. For cats, peak temporal frequencies for 36 single neurons from ref. 49 were combined with fitted peak temporal frequencies for 72 single neurons from ref. 52 to generate an aggregate distribution of peak temporal frequencies. For mice, fitted peak temporal frequencies for 192 single neurons from ref. 56 were combined with fitted peak temporal frequencies for 69 single neurons from ref. 57 to generate an aggregate distribution of peak temporal frequencies. This distribution was consistent with summary statistics reported in an additional mouse study using similar methods^55^. From the complete aggregate distributions for each animal, we measured the median and 25th and 75th quartiles.

### Cortical density

Cortical density per dva was estimated by taking the total number of V1 neurons and computing what the total 2D representation of neurons would be based on the surface area and depth of V1, and then dividing that by the total visual area in degrees. We estimated total numbers of neurons using the volume of V1 in mm^3^ times a density of 150,000 neurons/mm^3^ because this density has been a relatively consistent value reported in V1 for chimpanzees^104^, macaques^105^, marmosets^106^, and mice^107^. For humans, we used 1470 mm^2^ for surface area^108^ and 2.5 mm for depth^109^ to get 211 neurons/dva^2^. For marmosets, we used 200 mm^2^ for surface area^110^ and 1.1 mm for depth^111^ to get 28.7 neurons/dva^2^. For mice, we used 3 mm^2^ for surface area^112^ and 0.8 mm for depth^109^ to get 0.43 neurons/dva^2^.

### Statistics and reproducibility

All statistical tests were nonparametric based on the median and error bars were based on bootstrap analysis of the median by resampled data 1000 times, allowing repeats, to produce surrogate datasets of the same size. The 160th and 840th samples were used for the standard error of the median for all results. For data we collected on humans, marmosets, and mice, we present the entire distribution of all subjects grouped or single subject medians as single solid data points, an outline connecting the standard error of the median of grouped data across rate and size distributions, and an outline connecting the 25th and 75th quartiles of grouped data across rate and size distributions. For receptive field, temporal frequency and saccade data published previously by other authors, we present the median of grouped data or medians of single subjects as open circles and a dashed outline connecting the 25th and 75th quartiles of grouped data across y- and x-axis distributions. For statistical tests, bootstrapped data for one set of observations were used to find the percentile equal to the median of the compared set of observations. If the compared median was completely outside of the bootstrapped dataset, we reported that as *P* < 0.001.

## Data and code availability

The data presented in the figures and the model code used to generate the figures are available at Figshare: https://figshare.com/projects/Mammals_achieve_common_neural_coverage_of_visual_scenes_using_distinct_sampling_behaviors/136912

## Author Contributions

J.M.S., M.S., A.M., G.S.M., and N.J.P. designed research; J.M.S., M.S., C.B. and A.M. performed research; J.M.S., M.S., C.B. and A.M. analyzed data; and J.M.S. wrote the original draft; J.M.S., M.S., A.M., G.S.M., and N.J.P. edited the paper.

## Classification

Biological Sciences: Neuroscience

## Acknowledgments

We thank Allison Laudano for animal care and Jagruti Pattadkal for helping set up all marmoset experiments. We thank Theodore Weyand for sharing cat data and helpful discussions. This research was funded by ANR-20-NEUC-0002 (G.M.) and NIH-R01-NS120562 (N.P.).

## References

1 Yarbus, A. L. Eye Movements and Vision. (Springer US : Imprint: Springer, 1967).

2 Koch, C. & Ullman, S. Shifts in Selective Visual-Attention - Towards the Underlying Neural Circuitry. Hum Neurobiol 4, 219–227 (1985).

3 Hayhoe, M. & Ballard, D. Eye movements in natural behavior. Trends Cogn Sci 9, 188–194, doi:10.1016/j.tics.2005.02.009 (2005).

4 Najemnik, J. & Geisler, W. S. Optimal eye movement strategies in visual search. Nature 434, 387–391, doi:10.1038/nature03390 (2005).

5 Gegenfurtner, K. R. The Interaction Between Vision and Eye Movements (dagger). Perception 45, 1333–1357, doi:10.1177/0301006616657097 (2016).

6 Schutz, A. C., Braun, D. I. & Gegenfurtner, K. R. Eye movements and perception: a selective review. J Vis 11, doi:10.1167/11.5.9 (2011).

7 Land, M. F. Motion and vision: why animals move their eyes. J Comp Physiol A 185, 341–352, doi:DOI 10.1007/s003590050393 (1999).

8 Walls, G. L. The Evolutionary History of Eye Movements. Vision Res 2, 69–80, doi:Doi 10.1016/0042-6989(62)90064-0 (1962).

9 Land, M. Eye movements in man and other animals. Vision Res 162, 1–7, doi:10.1016/j.visres.2019.06.004 (2019).

10 Samonds, J. M., Geisler, W. S. & Priebe, N. J. Natural image and receptive field statistics predict saccade sizes. Nat Neurosci 21, 1591–1599, doi:10.1038/s41593-018-0255-5 (2018).

11 Goodchild, A. K., Ghosh, K. K. & Martin, P. R. Comparison of photoreceptor spatial density and ganglion cell morphology in the retina of human, macaque monkey, cat, and the marmoset Callithrix jacchus. J Comp Neurol 366, 55–75, doi:Doi 10.1002/(Sici)1096-9861(19960226)366:1<55::Aid-Cne5>3.0.Co;2-J (1996).

12 Szel, A. et al. Unique topographic separation of two spectral classes of cones in the mouse retina. J Comp Neurol 325, 327–342, doi:10.1002/cne.903250302 (1992).

13 Jeon, C. J., Strettoi, E. & Masland, R. H. The major cell populations of the mouse retina. J Neurosci 18, 8936–8946 (1998).

14 Rapaport, D. H. & Stone, J. The area centralis of the retina in the cat and other mammals: focal point for function and development of the visual system. Neuroscience 11, 289–301, doi:10.1016/0306-4522(84)90024-1 (1984).

15 Schiefer, U. et al. Angioscotoma detection with fundus-oriented perimetry. A study with dark and bright stimuli of different sizes. Vision Res 39, 1897–1909, doi:10.1016/s0042-6989(98)00295-8 (1999).

16 Wassle, H., Grunert, U., Rohrenbeck, J. & Boycott, B. B. Cortical Magnification Factor and the Ganglion-Cell Density of the Primate Retina. Nature 341, 643–646, doi:DOI 10.1038/341643a0 (1989).

17 Chaplin, T. A., Yu, H. H. & Rosa, M. G. P. Representation of the visual field in the primary visual area of the marmoset monkey: Magnification factors, point-image size, and proportionality to retinal ganglion cell density. J Comp Neurol 521, 1001–1019, doi:10.1002/cne.23215 (2013).

18 Albus, K. Quantitative Study of Projection Area of Central and Paracentral Visual-Field in Area-17 of Cat .2. Spatial Organization of Orientation Domain. Exp Brain Res 24, 181–202, doi:Doi 10.1007/Bf00234062 (1975).

19 Wilson, J. R. & Sherman, S. M. Receptive-Field Characteristics of Neurons in Cat Striate Cortex - Changes with Visual-Field Eccentricity. J Neurophysiol 39, 512–533 (1976).

20 Tusa, R. J., Palmer, L. A. & Rosenquist, A. C. Retinotopic Organization of Area-17 (Striate Cortex) in Cat. J Comp Neurol 177, 213–235, doi:DOI 10.1002/cne.901770204 (1978).

21 Van Essen, D. C., Newsome, W. T. & Maunsell, J. H. R. The Visual-Field Representation in Striate Cortex of the Macaque Monkey - Asymmetries, Anisotropies, and Individual Variability. Vision Res 24, 429–448, doi:Doi 10.1016/0042-6989(84)90041-5 (1984).

22 van Beest, E. H. et al. Mouse visual cortex contains a region of enhanced spatial resolution. Nat Commun 12, 4029, doi:10.1038/s41467-021-24311-5 (2021).

23 Tan, L., Ringach, D. L. & Trachtenberg, J. T. The Development of Receptive Field Tuning Properties in Mouse Binocular Primary Visual Cortex. J Neurosci 42, 3546–3556, doi:10.1523/JNEUROSCI.1702-21.2022 (2022).

24 Rhim, I., Coello-Reyes, G., Ko, H. K. & Nauhaus, I. Maps of cone opsin input to mouse V1 and higher visual areas. J Neurophysiol 117, 1674–1682, doi:10.1152/jn.00849.2016 (2017).

25 Adams, D. L. & Horton, J. C. Shadows cast by retinal blood vessels mapped in primary visual cortex. Science 298, 572–576, doi:10.1126/science.1074887 (2002).

26 Sprague, W. W., Cooper, E. A., Tosic, I. & Banks, M. S. Stereopsis is adaptive for the natural environment. Sci Adv 1, doi:ARTN e1400254 10.1126/sciadv.1400254 (2015).

27 Samonds, J. M., Choi, V. & Priebe, N. J. Mice Discriminate Stereoscopic Surfaces Without Fixating in Depth. J Neurosci 39, 8024–8037, doi:10.1523/Jneurosci.0895-19.2019 (2019).

28 La Chioma, A., Bonhoeffer, T. & Hubener, M. Area-Specific Mapping of Binocular Disparity across Mouse Visual Cortex. Curr Biol 29, 2954-+, doi:10.1016/j.cub.2019.07.037 (2019).

29 Melcher, D. & Colby, C. L. Trans-saccadic perception. Trends Cogn Sci 12, 466–473, doi:10.1016/j.tics.2008.09.003 (2008).

30 Cavanagh, P., Hunt, A. R., Afraz, A. & Rolfs, M. Visual stability based on remapping of attention pointers. Trends Cogn Sci 14, 147–153, doi:10.1016/j.tics.2010.01.007 (2010).

31 Gottlieb, J. From a different point of view: extrastriate cortex integrates information across saccades. Focus on "Remapping in human visual cortex". J Neurophysiol 97, 961–962, doi:10.1152/jn.01225.2006 (2007).

32 Ganmor, E., Landy, M. S. & Simoncelli, E. P. Near-optimal integration of orientation information across saccades. J Vis 15, 8, doi:10.1167/15.16.8 (2015).

33 Wolf, C. & Schutz, A. C. Trans-saccadic integration of peripheral and foveal feature information is close to optimal. J Vis 15, 1, doi:10.1167/15.16.1 (2015).

34 Stewart, E. E. M., Valsecchi, M. & Schutz, A. C. A review of interactions between peripheral and foveal vision. J Vis 20, 2, doi:10.1167/jov.20.12.2 (2020).

35 Muller, J. R., Metha, A. B., Krauskopf, J. & Lennie, P. Rapid adaptation in visual cortex to the structure of images. Science 285, 1405–1408, doi:DOI 10.1126/science.285.5432.1405 (1999).

36. Tring, E., Dipoppa, M. & Ringach, D. L. A power law of cortical adaptation. bioRxiv, doi:10.1101/2023.05.22.541834 (2023).

37 Sheth, S. A. et al. Linear and nonlinear relationships between neuronal activity, oxygen metabolism, and hemodynamic responses. Neuron 42, 347–355, doi:10.1016/s0896-6273(04)00221-1 (2004).

38 Priebe, N. J., Churchland, M. M. & Lisberger, S. G. Constraints on the source of short-term motion adaptation in macaque area MT. I. the role of input and intrinsic mechanisms. J Neurophysiol 88, 354–369, doi:10.1152/jn.00852.2001 (2002).

39 Patterson, C. A., Wissig, S. C. & Kohn, A. Distinct effects of brief and prolonged adaptation on orientation tuning in primary visual cortex. J Neurosci 33, 532–543, doi:10.1523/JNEUROSCI.3345-12.2013 (2013).

40 Jin, M., Beck, J. M. & Glickfeld, L. L. Neuronal Adaptation Reveals a Suboptimal Decoding of Orientation Tuned Populations in the Mouse Visual Cortex. J Neurosci 39, 3867–3881, doi:10.1523/JNEUROSCI.3172-18.2019 (2019).

41 Jin, M. & Glickfeld, L. L. Magnitude, time course, and specificity of rapid adaptation across mouse visual areas. J Neurophysiol 124, 245–258, doi:10.1152/jn.00758.2019 (2020).

42 Nelson, S. B. Temporal interactions in the cat visual system. I. Orientation-selective suppression in the visual cortex. J Neurosci 11, 344–356 (1991).

43 Mitchell, J. F., Reynolds, J. H. & Miller, C. T. Active Vision in Marmosets: A Model System for Visual Neuroscience. J Neurosci 34, 1183–1194, doi:10.1523/Jneurosci.3899-13.2014 (2014).

44 Otero-Millan, J., Macknik, S. L., Langston, R. E. & Martinez-Conde, S. An oculomotor continuum from exploration to fixation. P Natl Acad Sci USA 110, 6175–6180, doi:10.1073/pnas.1222715110 (2013).

45 Tatler, B. W., Baddeley, R. J. & Vincent, B. T. The long and the short of it: spatial statistics at fixation vary with saccade amplitude and task. Vision Res 46, 1857–1862, doi:10.1016/j.visres.2005.12.005 (2006).

46 Jansen, L., Onat, S. & Konig, P. Influence of disparity on fixation and saccades in free viewing of natural scenes. J Vis 9, 29 21–19, doi:10.1167/9.1.29 (2009).

47 Nikolaev, A. R., Jurica, P., Nakatani, C., Plomp, G. & van Leeuwen, C. Visual encoding and fixation target selection in free viewing: presaccadic brain potentials. Front Syst Neurosci 7, 26, doi:10.3389/fnsys.2013.00026 (2013).

48 Loh, Z., Hall, E. H., Cronin, D. & Henderson, J. M. Working memory control predicts fixation duration in scene-viewing. Psychol Res, doi:10.1007/s00426-022-01694-8 (2022).

49 Ikeda, H. & Wright, M. J. Spatial and Temporal Properties of ’Sustained’ and ’Transient’ Neurones in Area 17 of the Cat’s Visual Cortex. Exp Brain Res 22, 363–383 (1975).

50 Foster, K. H., Gaska, J. P., Nagler, M. & Pollen, D. A. Spatial and temporal frequency selectivity of neurones in visual cortical areas V1 and V2 of the macaque monkey. J Physiol 365, 331–363, doi:10.1113/jphysiol.1985.sp015776 (1985).

51 Hawken, M. J., Shapley, R. M. & Grosof, D. H. Temporal-frequency selectivity in monkey visual cortex. Vis Neurosci 13, 477–492, doi:10.1017/s0952523800008154 (1996).

52 Allison, J. D., Smith, K. R. & Bonds, A. B. Temporal-frequency tuning of cross-orientation suppression in the cat striate cortex. Vis Neurosci 18, 941–948 (2001).

53 Webb, B. S., Dhruv, N. T., Solomon, S. G., Tailby, C. & Lennie, P. Early and late mechanisms of surround suppression in striate cortex of macaque. J Neurosci 25, 11666–11675, doi:10.1523/JNEUROSCI.3414-05.2005 (2005).

54 Priebe, N. J., Lisberger, S. G. & Movshon, J. A. Tuning for spatiotemporal frequency and speed in directionally selective neurons of macaque striate cortex. J Neurosci 26, 2941–2950, doi:10.1523/JNEUROSCI.3936-05.2006 (2006).

55 Niell, C. M. & Stryker, M. P. Highly selective receptive fields in mouse visual cortex. J Neurosci 28, 7520–7536, doi:10.1523/Jneurosci.0623-08.2008 (2008).

56 Gao, E. Q., DeAngelis, G. C. & Burkhalter, A. Parallel Input Channels to Mouse Primary Visual Cortex. J Neurosci 30, 5912–5926, doi:10.1523/Jneurosci.6456-09.2010 (2010).

57 Van den Bergh, G., Zhang, B., Arckens, L. & Chino, Y. M. Receptive-field Properties of V1 and V2 Neurons in Mice and Macaque Monkeys. J Comp Neurol 518, 2051–2070, doi:10.1002/cne.22321 (2010).

58 Yu, H. H. et al. Spatial and temporal frequency tuning in striate cortex: functional uniformity and specializations related to receptive field eccentricity. Eur J Neurosci 31, 1043–1062, doi:10.1111/j.1460-9568.2010.07118.x (2010).

59 Durand, S. et al. A Comparison of Visual Response Properties in the Lateral Geniculate Nucleus and Primary Visual Cortex of Awake and Anesthetized Mice. J Neurosci 36, 12144–12156, doi:10.1523/JNEUROSCI.1741-16.2016 (2016).

60 Alitto, H. J. & Usrey, W. M. Influence of contrast on orientation and temporal frequency tuning in ferret primary visual cortex. J Neurophysiol 91, 2797–2808, doi:10.1152/jn.00943.2003 (2004).

61 Camillo, D., Ahmadlou, M. & Heimel, J. A. Contrast-Dependence of Temporal Frequency Tuning in Mouse V1. Front Neurosci 14, 868, doi:10.3389/fnins.2020.00868 (2020).

62 Seiple, W., Rosen, R. B. & Garcia, P. M. Abnormal fixation in individuals with age-related macular degeneration when viewing an image of a face. Optom Vis Sci 90, 45–56, doi:10.1097/OPX.0b013e3182794775 (2013).

63 Kwon, M. Y., Nandy, A. S. & Tjan, B. S. Rapid and Persistent Adaptability of Human Oculomotor Control in Response to Simulated Central Vision Loss. Curr Biol 23, 1663–1669, doi:10.1016/j.cub.2013.06.056 (2013).

64 Chen, D., Otero-Millan, J., Kumar, P., Shaikh, A. G. & Ghasia, F. F. Visual Search in Amblyopia: Abnormal Fixational Eye Movements and Suboptimal Sampling Strategies. Invest Ophthalmol Vis Sci 59, 4506–4517, doi:10.1167/iovs.18-24794 (2018).

65 Tang, S. et al. Effects of visual blur on microsaccades during visual exploration. J Eye Mov Res 12, doi:10.16910/jemr.12.6.10 (2019).

66 Jang, J., Song, M. & Paik, S. B. Retino-Cortical Mapping Ratio Predicts Columnar and Salt-and-Pepper Organization in Mammalian Visual Cortex. Cell Rep 30, 3270–3279 e3273, doi:10.1016/j.celrep.2020.02.038 (2020).

67 Fasoli, D. & Panzeri, S. Mathematical studies of the dynamics of finite-size binary neural networks: A review of recent progress. Math Biosci Eng 16, 8025–8059, doi:10.3934/mbe.2019404 (2019).

68 Meyer, A. F., O’Keefe, J. & Poort, J. Two Distinct Types of Eye-Head Coupling in Freely Moving Mice. Curr Biol 30, 2116–2130 e2116, doi:10.1016/j.cub.2020.04.042 (2020).

69 Michaiel, A. M., Abe, E. T. & Niell, C. M. Dynamics of gaze control during prey capture in freely moving mice. Elife 9, doi:10.7554/eLife.57458 (2020).

70 Zahler, S. H., Taylor, D. E., Wong, J. Y., Adams, J. M. & Feinberg, E. H. Superior colliculus drives stimulus-evoked directionally biased saccades and attempted head movements in head-fixed mice. Elife 10, doi:10.7554/eLife.73081 (2021).

71 Harwood, M. R., Madelain, L., Krauzlis, R. J. & Wallman, J. The spatial scale of attention strongly modulates saccade latencies. J Neurophysiol 99, 1743–1757, doi:10.1152/jn.00589.2007 (2008).

72 De Vries, J. P., Azadi, R. & Harwood, M. R. The saccadic size-latency phenomenon explored: Proximal target size is a determining factor in the saccade latency. Vision Res 129, 87–97, doi:10.1016/j.visres.2016.09.006 (2016).

73 Poletti, M., Intoy, J. & Rucci, M. Accuracy and precision of small saccades. Sci Rep 10, 16097, doi:10.1038/s41598-020-72432-6 (2020).

74 Segal, I. Y. et al. Decorrelation of retinal response to natural scenes by fixational eye movements. P Natl Acad Sci USA 112, 3110–3115, doi:10.1073/pnas.1412059112 (2015).

75 Kuang, X. T., Poletti, M., Victor, J. D. & Rucci, M. Temporal Encoding of Spatial Information during Active Visual Fixation. Curr Biol 22, 510–514, doi:10.1016/j.cub.2012.01.050 (2012).

76 Mostofi, N. et al. Spatiotemporal Content of Saccade Transients. Curr Biol 30, 3999–4008 e3992, doi:10.1016/j.cub.2020.07.085 (2020).

77. Parker, P. R., et al. A dynamic sequence of visual processing initiated by gaze shifts. BioRxiv, doi:10.1101/2022.08.23.504847 (2022).

78 Castet, E. & Masson, G. S. Motion perception during saccadic eye movements. Nat Neurosci 3, 177–183, doi:10.1038/72124 (2000).

79 Castet, E., Jeanjean, S. & Masson, G. S. Motion perception of saccade-induced retinal translation. Proc Natl Acad Sci U S A 99, 15159–15163, doi:10.1073/pnas.232377199 (2002).

80 Miles, F. A., Kawano, K. & Optican, L. M. Short-latency ocular following responses of monkey. I. Dependence on temporospatial properties of visual input. J Neurophysiol 56, 1321–1354, doi:10.1152/jn.1986.56.5.1321 (1986).

81 Schweitzer, R. & Rolfs, M. Intra-saccadic motion streaks as cues to linking object locations across saccades. J Vis 20, 17, doi:10.1167/jov.20.4.17 (2020).

82 Schweitzer, R. & Rolfs, M. Intrasaccadic motion streaks jump-start gaze correction. Sci Adv 7, doi:10.1126/sciadv.abf2218 (2021).

83 Rolfs, M. & Schweitzer, R. Coupling perception to action through incidental sensory consequences of motor behaviour. Nature Reviews Psychology 1, 112–123 (2022).

84 Johnson, K. P. et al. Cell-type-specific binocular vision guides predation in mice. Neuron 109, 1527–1539 e1524, doi:10.1016/j.neuron.2021.03.010 (2021).

85 Groner, M. T., Groner, R. & von Muhlenen, A. The effect of spatial frequency content on parameters of eye movements. Psychol Res-Psych Fo 72, 601–608, doi:10.1007/s00426-008-0167-1 (2008).

86 Intoy, J. & Rucci, M. Finely tuned eye movements enhance visual acuity. Nat Commun 11, 795, doi:10.1038/s41467-020-14616-2 (2020).

87 Niemeier, M., Crawford, J. D. & Tweed, D. B. Optimal transsaccadic integration explains distorted spatial perception. Nature 422, 76–80, doi:10.1038/nature01439 (2003).

88 Oostwoud Wijdenes, L., Marshall, L. & Bays, P. M. Evidence for Optimal Integration of Visual Feature Representations across Saccades. J Neurosci 35, 10146–10153, doi:10.1523/JNEUROSCI.1040-15.2015 (2015).

89 Henderson, J. M. & Hollingworth, A. Eye movements and visual memory: detecting changes to saccade targets in scenes. Percept Psychophys 65, 58–71, doi:10.3758/bf03194783 (2003).

90 Scholl, B., Pattadkal, J. J., Dilly, G. A., Priebe, N. J. & Zemelman, B. V. Local Integration Accounts for Weak Selectivity of Mouse Neocortical Parvalbumin Interneurons. Neuron 87, 424–436, doi:10.1016/j.neuron.2015.06.030 (2015).

91 Choi, V. & Priebe, N. J. Interocular velocity cues elicit vergence eye movements in mice. J Neurophysiol 124, 623–633, doi:10.1152/jn.00697.2019 (2020).

92 Pelli, D. G. The VideoToolbox software for visual psychophysics: transforming numbers into movies. Spat Vis 10, 437–442 (1997).

93 Brainard, D. H. The psychophysics toolbox. Spatial Vision 10, 433–436, doi:Doi 10.1163/156856897X00357 (1997).

94 Cornelissen, F. W., Peters, E. M. & Palmer, J. The Eyelink Toolbox: eye tracking with MATLAB and the Psychophysics Toolbox. Behav Res Methods Instrum Comput 34, 613–617, doi:10.3758/bf03195489 (2002).

95 Olmos, A. & Kingdom, F. A. A. A biologically inspired algorithm for the recovery of shading and reflectance images. Perception 33, 1463–1473, doi:10.1068/p5321 (2004).

96 Mathis, A. et al. DeepLabCut: markerless pose estimation of user-defined body parts with deep learning. Nat Neurosci 21, 1281–1289, doi:10.1038/s41593-018-0209-y (2018).

97 Mitchell, J. F., Priebe, N. J. & Miller, C. T. Motion dependence of smooth pursuit eye movements in the marmoset. J Neurophysiol 113, 3954–3960, doi:10.1152/jn.00197.2015 (2015).

98 Menzel, C. R. & Menzel, E. W. Head-Cocking and Visual Exploration in Marmosets (Sanguinus fuscicollis). Behaviour 75, 219–234 (1980).

99 Umino, Y., Pasquale, R. & Solessio, E. Visual Temporal Contrast Sensitivity in the Behaving Mouse Shares Fundamental Properties with Human Psychophysics. Eneuro 5, doi:10.1523/ENEURO.0181-18.2018 (2018).

100 Scholl, B., Pattadkal, J. J. & Priebe, N. J. Binocular Disparity Selectivity Weakened after Monocular Deprivation in Mouse V1. J Neurosci 37, 6517–6526, doi:10.1523/Jneurosci.1193-16.2017 (2017).

101 Lee, D. & Malpeli, J. G. Effects of saccades on the activity of neurons in the cat lateral geniculate nucleus. J Neurophysiol 79, 922–936 (1998).

102 Kording, K. P., Kayser, C., Betsch, B. Y. & Konig, P. Non-contact eye-tracking on cats. J Neurosci Meth 110, 103–111, doi:Doi 10.1016/S0165-0270(01)00423-X (2001).

103 Roth, M. M. et al. Thalamic nuclei convey diverse contextual information to layer 1 of visual cortex. Nat Neurosci 19, 299-+, doi:10.1038/nn.4197 (2016).

104 Miller, D. J., Balaram, P., Young, N. A. & Kaas, J. H. Three counting methods agree on cell and neuron number in chimpanzee primary visual cortex. Front Neuroanat 8, 36, doi:10.3389/fnana.2014.00036 (2014).

105 O’Kusky, J. & Colonnier, M. A laminar analysis of the number of neurons, glia, and synapses in the adult cortex (area 17) of adult macaque monkeys. J Comp Neurol 210, 278–290, doi:10.1002/cne.902100307 (1982).

106 Atapour, N. et al. Neuronal Distribution Across the Cerebral Cortex of the Marmoset Monkey (Callithrix jacchus). Cereb Cortex 29, 3836–3863, doi:10.1093/cercor/bhy263 (2019).

107 Keller, D., Ero, C. & Markram, H. Cell Densities in the Mouse Brain: A Systematic Review. Front Neuroanat 12, 83, doi:10.3389/fnana.2018.00083 (2018).

108 Dougherty, R. F. et al. Visual field representations and locations of visual areas V1/2/3 in human visual cortex. J Vis 3, 586–598, doi:10.1167/3.10.1 (2003).

109 DeFelipe, J. The evolution of the brain, the human nature of cortical circuits, and intellectual creativity. Front Neuroanat 5, doi:ARTN 29 10.3389/fnana.2011.00029 (2011).

110 Solomon, S. G. & Rosa, M. G. A simpler primate brain: the visual system of the marmoset monkey. Front Neural Circuits 8, 96, doi:10.3389/fncir.2014.00096 (2014).

111 Balaram, P. & Kaas, J. H. Towards a unified scheme of cortical lamination for primary visual cortex across primates: insights from NeuN and VGLUT2 immunoreactivity. Front Neuroanat 8, 81, doi:10.3389/fnana.2014.00081 (2014).

112 Garrett, M. E., Nauhaus, I., Marshel, J. H. & Callaway, E. M. Topography and areal organization of mouse visual cortex. J Neurosci 34, 12587–12600, doi:10.1523/JNEUROSCI.1124-14.2014 (2014).

