## Supplemental Fig 6 for "Mammals achieve common neural coverage of visual scenes using distinct sampling behaviors"

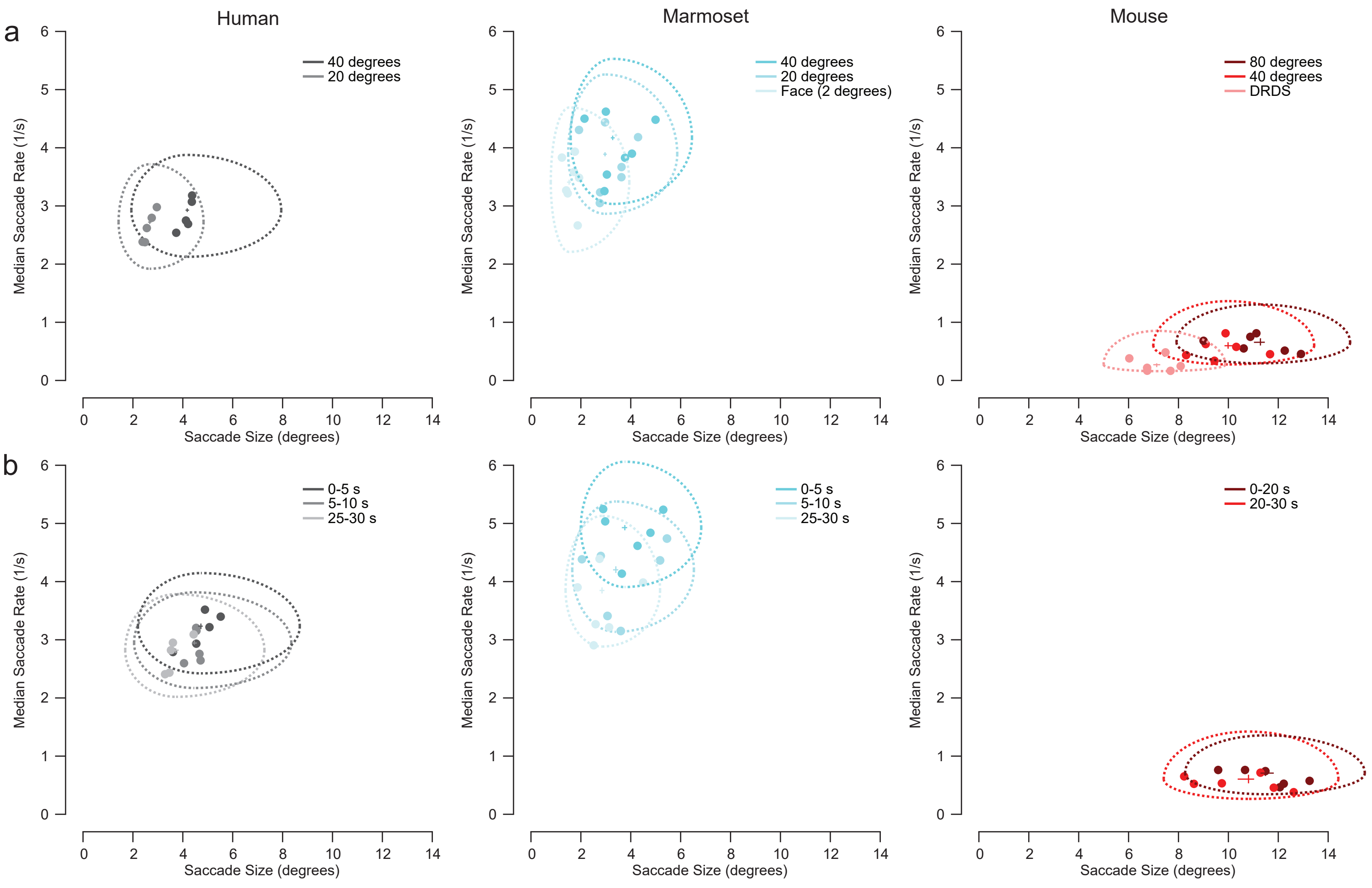


**Supplemental Figure 6. Saccade size and rate decrease with decreasing image size and novelty**
**a**, Saccade rate versus size distributions for small and large images for humans, marmosets, and mice (same data as Fig. 1f and Supplemental Fig. 2). Lightest colors represent data for n = 7 marmosets performing fixation training (n = 8,738 saccades) and n = 6 mice discriminating DRDS disparity (n = 2,016 saccades). Each solid point represents the medians of a single subject. The large outline represents 25th and 75th percentiles and the small cross represents the standard error of the median for the distribution of all subjects. Due to the large number of samples, the standard error of the median outlines are smaller than even the data points. For all species, the saccade size and rate were substantially and significantly smaller for these tasks compared to saccades generated when viewing natural images (bootstrapped, $p < 0.001$ for all comparisons).
**b**, Image size is not the only dimension that factors into the amount of sampled visual information over time. Visual information also becomes redundant and less informative over time, if there are no changes or updates in visual information, as for instance with static images. We measured changes in saccade rate and size over the entire 30 s presentations for humans, marmosets, and mice. Saccade rate versus size over time for n = 5 humans (n = 10,172; n = 8,949; and n = 7,500 saccades), n = 7 marmosets (n = 6,070; n = 5,284; and n = 4,173 saccades), and n = 6 mice (n = 947 and n = 408 saccades) viewing large images. For humans and marmosets, saccade rates and sizes quickly decreased over time (within the first 5 s of viewing an image; bootstrapped, $p < 0.001$ for all comparisons). This decrease was quicker for animals that saccade at faster rates. Marmosets had a quicker decrease in rates and sizes compared to humans (bootstrapped, 14.7 vs 8.9% and 9.6 vs 5.4% decrease in the first 5 s, $p < 0.001$ and $p = 0.008$, respectively) and there was only a noticeable decrease in saccade rates and sizes for mice after viewing an image for 20 s (bootstrapped, $p = 0.06$ and $p = 0.003$, respectively).
