## Supplemental Fig 3 for "Mammals achieve common neural coverage of visual scenes using distinct sampling behaviors"

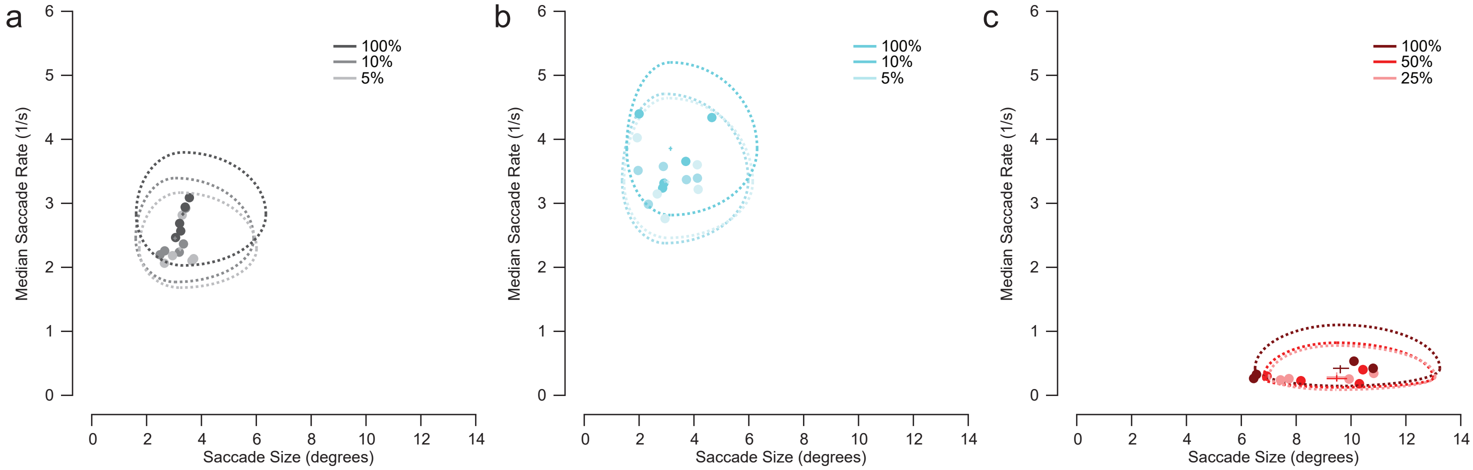


**Supplemental Figure 3. Saccade rates decrease with decreasing contrast**
**a**, Each solid point represents the medians of a single subject. The large outline represents 25th and 75th percentiles and the small cross represents the standard error of the median for the distribution of all subjects. Due to the large number of samples, the standard error of the median outlines are smaller than even the data points. Saccade rate versus size distributions for n = 5 humans for images with progressively reduced contrast (small and large images combined). (100%: n = 92,430 saccades; 10%: n = 42,767 saccades; 5%: n = 41,639 saccades)
**b**, Same data for n = 5 marmosets. (100%: n = 40,474 saccades; 10%: n = 31,927 saccades; 5%: n = 37,294 saccades)
**c**, Same data for n = 4 mice. (100%: n = 472 saccades; 50%: n = 338 saccades; 25%: n = 274 saccades)
For all three animals, there was a clear decrease in saccade rate for the images with reduced contrast (from dark to light colors; bootstrapped, 5 and 10 versus 100%, $p < 0.001$ for all comparisons for humans and marmosets; 25 and 50 versus 100%, $p = 0.008$ and 0.01, respectively for mice).
