## Supplemental Fig 2 for "Mammals achieve common neural coverage of visual scenes using distinct sampling behaviors"

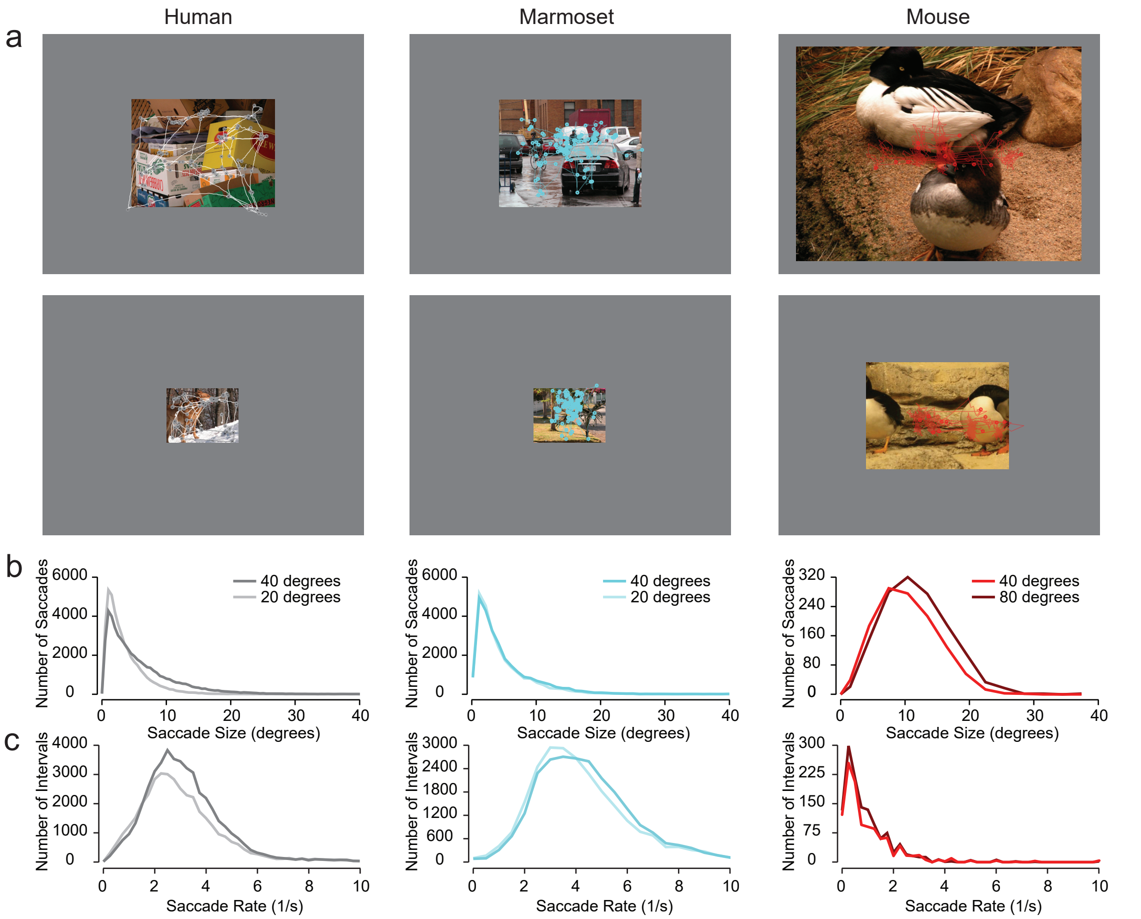


**Supplemental Figure 2. Saccadic eye movement statistics while viewing natural images**
**a**, Eye traces and fixation points (circles) for humans, marmosets and mice viewing an example large (top row) or small (bottom row) image for 30 s. For mice, we plotted fixation points with respect to an assumed projection of binocular photoreceptors by aligning the center of the image with the median eye position, since standard fixation calibration procedures are not possible.
**b**, Histograms of saccade sizes for N = 5 humans each viewing 80 large (n = 49,907 saccades) and small (n = 42,523 saccades) images, N = 7 marmosets each viewing 24 large (n = 25,531 saccades) and small images (n = 24,781 saccades), and N = 6 mice each viewing 8-18 large (n = 1,355 saccades) and small (n = 1,145 saccades) images[10].
**c**, Histograms of saccade rates (1/intersaccadic interval) for the same subjects.
