## Supplemental Fig 1 for "Mammals achieve common neural coverage of visual scenes using distinct sampling behaviors"

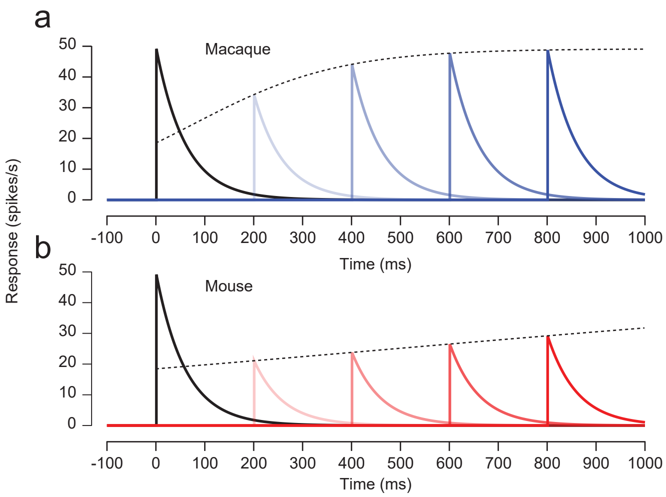


**Supplemental Figure 1. Example V1 responses modeled with a divisive gain time constant**
**a**, Example model macaque V1 responses to two repeated presentations of the same stimulus. Black represents neural activity to the first presentation and different shades of blue depict the neural activity after the second presentation in progressively more delayed conditions.
**b**, Same format as **a** for mouse model V1 responses.
