## Supplemental Fig 7 for "Mammals achieve common neural coverage of visual scenes using distinct sampling behaviors"

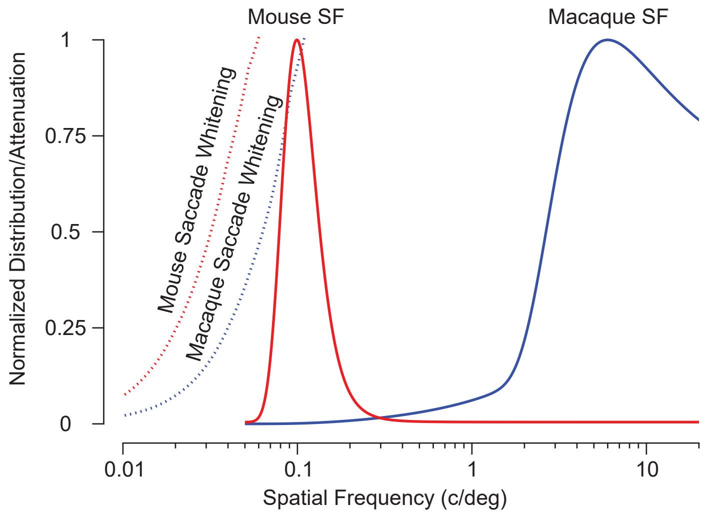


**Supplemental Figure 7. Saccade and fixation dynamics whiten natural inputs to V1**
Plots of spatial frequency distributions of model V1 receptive fields for mice (solid red) and macaques (solid blue) used to predict saccade sizes based on decorrelation of responses[2] compared to the attenuation of spatial frequencies relative to natural images caused by >7-degree saccades (dashed red) and 3-4-degree saccades (dashed blue). We estimated attenuation by dividing the spectral data from Fig. 4c by the spectral density of natural images in ref. 57. Larger saccades and lower temporal frequencies shift the attenuation curve to lower spatial frequencies, while smaller saccades and higher temporal frequencies shift the attenuation to higher spatial frequencies
