## Supplemental Fig 5 for "Mammals achieve common neural coverage of visual scenes using distinct sampling behaviors"

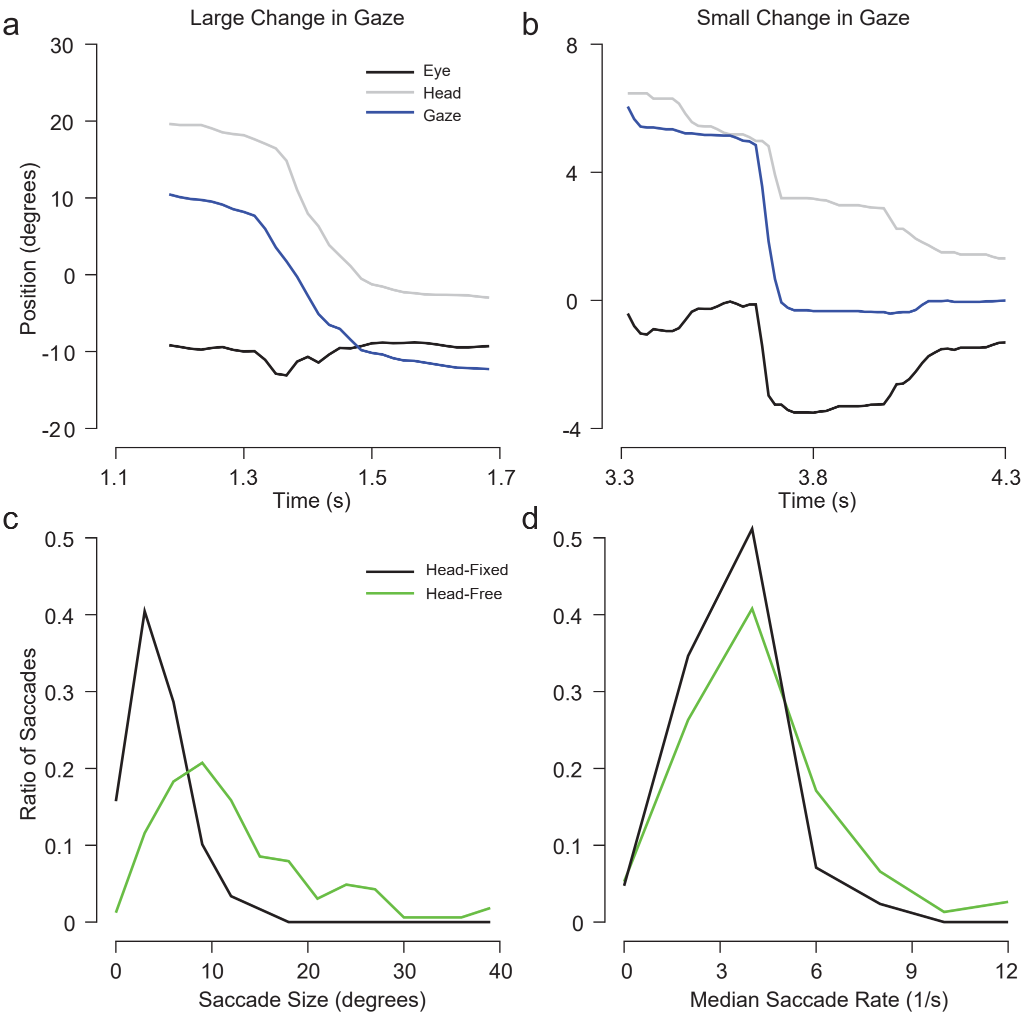


**Supplemental Figure 5. Contribution of head and eye movements to marmoset gaze changes**
**a**, An example of a large change in gaze that is mostly driven by a saccadic head movement. Vestibular ocular reflex (VOR) movements at the beginning and end of the head movement lead to an overall faster gaze changes separated by two stable periods of fixation.
**b**, An example of a small change in gaze. Although most of the movement is the result of the eyes, there is still a significant contributing saccadic head movement and there are similar VOR dynamics as those observed with a large change in gaze.
**c**, Distributions of saccadic gaze change sizes for head fixed and freely moving (head-free) marmosets.
**d**, Distributions of saccadic gaze change rates for head fixed and head-free marmosets.
