## Supplemental Fig 4 for "Mammals achieve common neural coverage of visual scenes using distinct sampling behaviors"

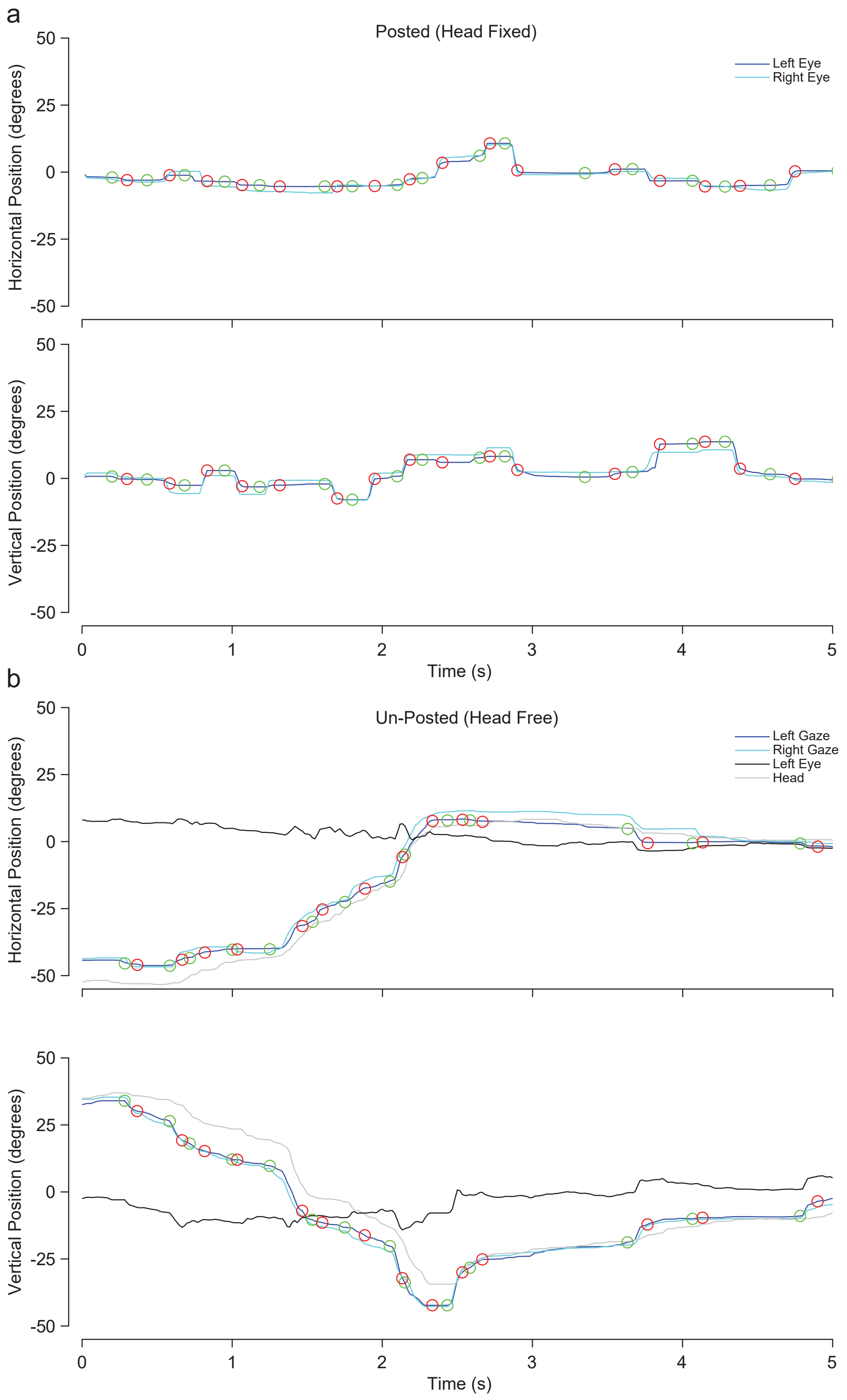


**Supplemental Figure 4. Gaze position over time for posted and unposted marmosets**
**a**, Horizontal and vertical pupil position for both eyes for a head-fixed marmoset. Green circles are saccade onset and red circles are saccade offset.
**b**, Same gaze data for the same marmoset now able to move their head freely unposted (head-free). Forehead position and left eye position relative to the head are included with the gaze traces. Note that the eyes never rotate out more than ±15 degrees and head movements drive the gaze changes.
